# Molecular identification of the *Trypanosoma* (*Herpetosoma*) *lewisi* clade in black rats (*Rattus rattus*) from Australia

**DOI:** 10.1101/819060

**Authors:** Siobhon L. Egan, Casey L. Taylor, Jill M. Austen, Peter B. Banks, Liisa A. Ahlstrom, Una M. Ryan, Peter J. Irwin, Charlotte L. Oskam

**Affiliations:** Vector and Waterborne Pathogens Research Group, College of Science, Health, Engineering and Education, Murdoch University, Perth, Western Australia, Australia; School of Life and Environmental Sciences, The University of Sydney, Sydney, New South Wales, Australia; Bayer Australia Ltd, Animal Health, Pymble, New South Wales, Australia

**Keywords:** *Trypanosoma lewisi*, *Rattus rattus*, Australia, black rats, ship rats

## Abstract

Invasive rodent species are known hosts for a diverse range of infectious microorganisms and have long been associated with the spread of disease globally. The present study describes molecular evidence for the presence of a *Trypanosoma* sp. from black rats (*Rattus rattus*) in northern Sydney, Australia. Sequences of the 18S ribosomal RNA (rRNA) locus were obtained in two out of eleven (18%) blood samples with subsequent phylogenetic analysis confirming the identity within the *Trypanosoma lewisi* clade.

## Introduction

Black rats (*Rattus rattus*) are distributed throughout the world and considered one of the most significant invasive species. In Australia, black rats became established alongside European settlement during the 1770’s, although the precise date of their first arrival on the continent is unclear (Banks and Hughes 2012). Black rats can act as amplifying hosts for a diverse range of pathogens that can affect humans, wildlife and domestic animals and a recent review of black rats in Europe identified at least 20 zoonotic infectious agents associated with the species (Strand and Lundkvist 2019). However, despite the global recognition of these rodents as hosts of pathogens, there is a relatively limited understanding of the range of infectious agents present in Australian populations of black rats (Banks and Hughes 2012).

Trypanosomes are a group of flagellate protozoan parasites, the vast majority of which are transmitted by blood-feeding invertebrates. Worldwide at least 44 trypanosome species are known to infect rodents (Dybing et al. 2016). In Australia, recent research has revealed the presence of several novel trypanosomes infecting native Australian marsupials (Thompson et al. 2014), however investigation into the presence of trypanosomes in Australian rodents, either native or introduced, has been lacking in recent years.

The results shared in this short communication form part of a broader ongoing investigation into vector-borne microorganisms present in Australia. To the authors’ knowledge, this study provides the first molecular identification of *Trypanosoma lewisi-*like organisms from black rats on mainland Australia.

## Methods

Small mammal trapping was conducted during April and May 2019 at two sites in northern Sydney, Australia; Irrawong Reserve and Warriewood Wetlands, Warriewood (−31.69°, 151.28°) and North Head, Manly (−33.81°, 151.29°). Two transects of 20 trap stations were set up at each site, with each station including one Elliot type B trap (46 × 15.5 × 15 cm) and one medium sized cage trap (72 × 32 × 31 cm) to target small and medium sized mammals. Traps were baited with peanut butter and oat balls and set for 3 consecutive nights. The sampling was conducted with approval of the Animal Ethics Committees of Murdoch University (Permit number R3026/18) and the University of Sydney (Permit number 2018/1429). Venous blood was collected into 1mL EDTA tubes for the detection of haemoparasites. Thin blood smears were prepared and stained with modified Wright-Giemsa. Blood films were inspected by light microscopy (Olympus BX51) for the presence of trypanosomes at x 400 magnification and under oil immersion (x 1000). Total genomic DNA was extracted from 200 μl of blood using a MasterPure DNA purification kit (Epicentre^®^ Biotechnologies, Madison, Wisconsin, U.S.A) following the manufacturer’s recommendations. Where 200 μl of blood was not available, PBS was used to make samples up to 200 μl. DNA was eluted in 30 μl of TE buffer and stored at −20°C.

Blood samples were screened for the presence of *Trypanosoma* spp. using a nested PCR approach targeting a ~550 bp product of the 18S ribosomal RNA (rRNA) gene with external primers TRY927F/TRY927R and internal primers SSU561F/SSU561R, as previously described (Noyes et al. 1999). Reactions were carried out in 25 μl volumes, 2 μl of gDNA was added to the primary PCR and 1 μl of the primary product was used as a template for the secondary assay. PCR products were electrophoresed on a 1% agarose gel stained with SYBR safe (Invitrogen, USA), and amplicons of the correct size were excised and purified using previously described methods (Yang et al. 2013). Sanger sequencing was carried out using internal primer sets in both directions and sequencing was performed at the Australian Genome Research Facility (Perth, Australia). Samples that returned a positive identification for *Trypanosoma lewisi*-like were further investigated. A near full-length fragment of the 18S rRNA locus was obtained using two nested PCR assays. Reactions were carried out in 25 μl volumes using external primers SLF/S762 and internal primer sets S823/S662 and S825/ SLIR as described (McInnes et al. 2009). Gel electrophoresis and Sanger sequencing using internal primers in both directions were carried out as above. No-template and extraction controls were included throughout the laboratory processes. Extractions, pre-PCR and post-PCR procedures were performed in laboratories physically separated from each other in order to minimise the risk of contamination. In addition, no *T. lewisi* species have been previously isolated or amplified in the specific laboratories used.

Nucleotide sequences from *Trypanosoma* species were retrieved from GenBank (Benson et al. 2017) and aligned with sequences obtained in the present study using MUSCLE (Edgar 2004), gaps were removed using Gblocks (Castresana 2000) with less stringent parameters. The final alignments were imported into MEGA 7 (Kumar et al. 2016), and the most appropriate nucleotide selection model was selected using the dedicated feature based on the Bayesian Information Criterion (BIC). Bayesian phylogenetic reconstruction was conducted in MrBayes v3.2.6 (Ronquist et al. 2012) using a MCMC length of 1,100,00, burn in of 10,000 and sub-sampling every 200 iterations. Genetic distances were calculated using the Kimura model, positions containing gaps and missing data were eliminated.

## Results and Discussion

In total, 11 black rat blood samples were collected for analysis from Warriewood Wetlands (n=4), and North Head (n=7). Two rat samples from North Head were positive for *Trypanosoma* species by molecular methods, and of these a blood smear was only available in one case. Unfortunately, no trypomastigote stages were observed by light microscopy despite prolonged searching of the cell layer. Black rat samples that were negative for molecular evidence of trypanosomes were also screened by microscopy, however this also did not return any positive observations. The absence of a morphological identification in this report is disappointing, however it is not unexpected when parasites reside in their natural host. Mackerras (1959) reported that rats (*R. rattus*) experimentally infected with *T. lewisi* go through an acute phase where parasites multiply rapidly, followed by a chronic phase, during which parasite numbers progressively diminish and disappear from circulation.

Initial screening produced ~550 bp product of the 18S rRNA gene in samples BR042 and BR048, these sequences were 100% identical to each other. A near full length 18S rRNA sequence (1,928 bp) was obtained from both samples also confirming that the sequences were 100% identical and a representative sequence of the 18S rRNA gene from sample BR042 was used for phylogenetic purposes (GenBank accession MN512227).

Phylogenetic analysis of the shorter (326 bp) 18S rRNA gene alignment was used in order to include a wider variety of reference sequences, in particular for the context of the present study to include the only other *T. lewisi*-like sequences from Australia (Averis et al. 2009). Figure 1 shows the phylogeny of the *Trypanosoma* genus (Fig 1a) and the resolution within the *T. lewisi* clade (Fig 1b). As demonstrated by the polytomy present in Fig 1b, this short region of the 18S rRNA gene is insufficient in the differentiation of members within the *T. lewisi* clade. Due to the speed at which the 18S rRNA locus has evolved, short regions of this locus have been reported as being unsuitable for inference of evolutionary relationships between *Trypanosoma* species (Hamilton and Stevens, 2011).

**Fig 1.**
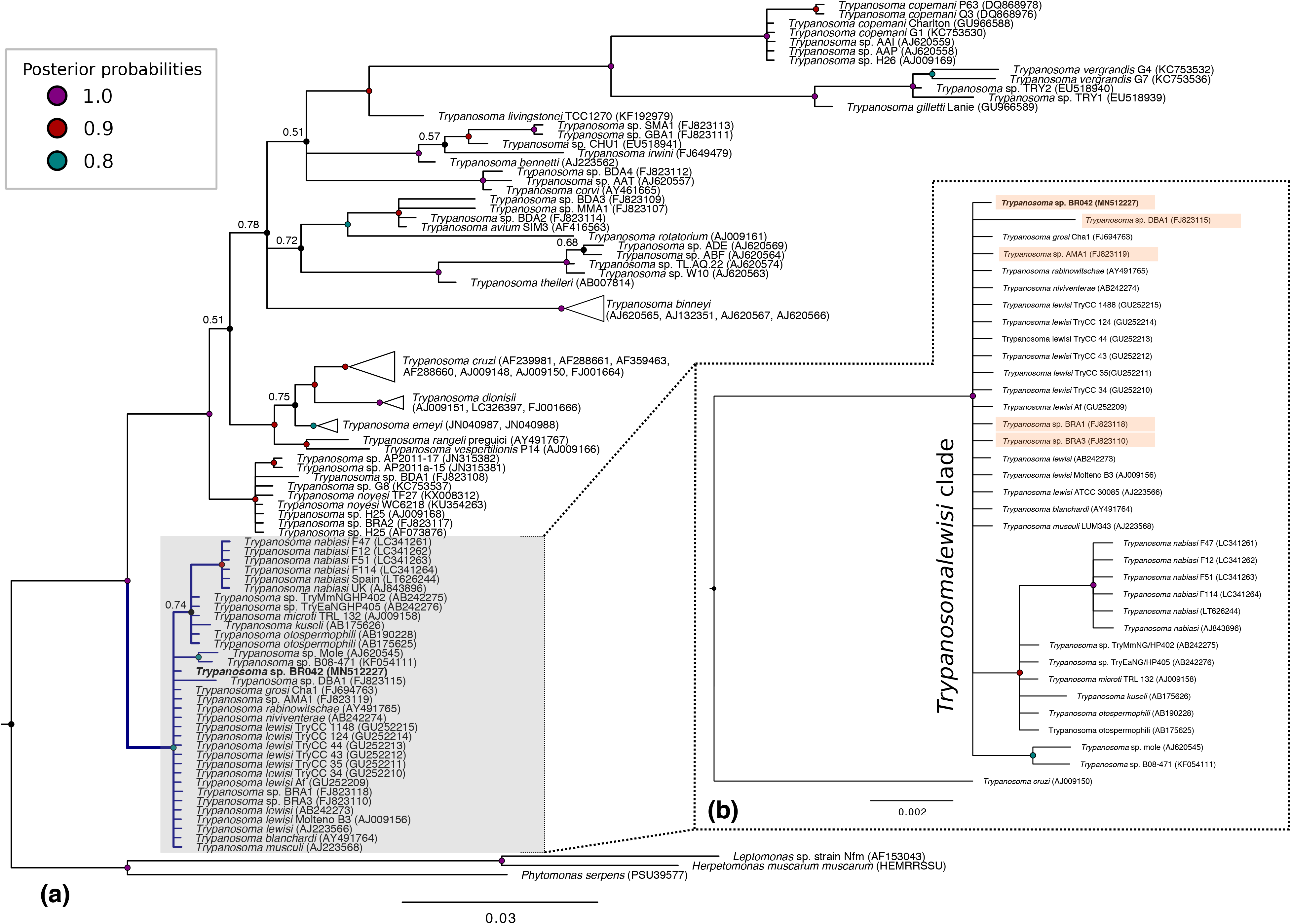
Bayesian phylogenetic reconstruction of *Trypanosoma* based on a 326 bp fragment of the 18S rRNA locus using the HKY + G substitution model. All positions containing gaps and missing data were eliminated. Phylogeny of the *Trypanosoma* genus (a) and insert tree (b) to show resolution of the *T. lewisi*-clade based on a short 326 bp to include a larger set of reference sequences, sequences obtained from Australia are shaded. All posterior probabilities at branch nodes >0.8 (see colour key on figure) unless indicated, number of substitutions per nucleotide position is represented by the scale bar. Collapsed nodes represented with triangular branches. New sequence from the present study is designated in bold.

Reconstruction of phylogenetic relationships over a longer region (1,627 bp) of the 18S rRNA gene exhibited superior resolution within the *T. lewisi* clade (Fig 2). In this phylogeny, sequences obtained from Australian black rats in the present study did not fall within the *T. lewisi* sensu stricto clade; instead they formed a distinct group that branched separately from other reference sequences. Pairwise distance analysis over a 1,627 bp alignment of the 18S rRNA gene demonstrated sequences from the black rat were 99.5% similar to *Trypanosoma microti* (AJ009158). The next most similar sequences were *Trypanosoma* sequences from voles in Japan (AB242275, AB242276) and a flea from Czech Republic (KF054111), all of which were 99.4% similar. Members of the *T. lewisi* sensu stricto clade, as shown in Fig 2., were all 100% identical to each other over the 1,627 bp alignment. These were the third most similar sequences (99.3%) to the *Trypanosoma* sp. identified in the present study. The phylogeny in the present study supports previous research by Hamilton et al. (2005) showing that the *T. lewisi* clade can be divided into two subclades, consisting of *T. lewisi*, *T. musculi*, *T. rabinowitschae*, *T. blanchardi* and *T. grosi* in subclade one and *T. nabiasi*, *T. microti, T. otospermophili* in subclade two. Sequences obtained in the present study from Australian black rats fall within subclade two of the *T. lewisi* clade.

**Fig 2.**
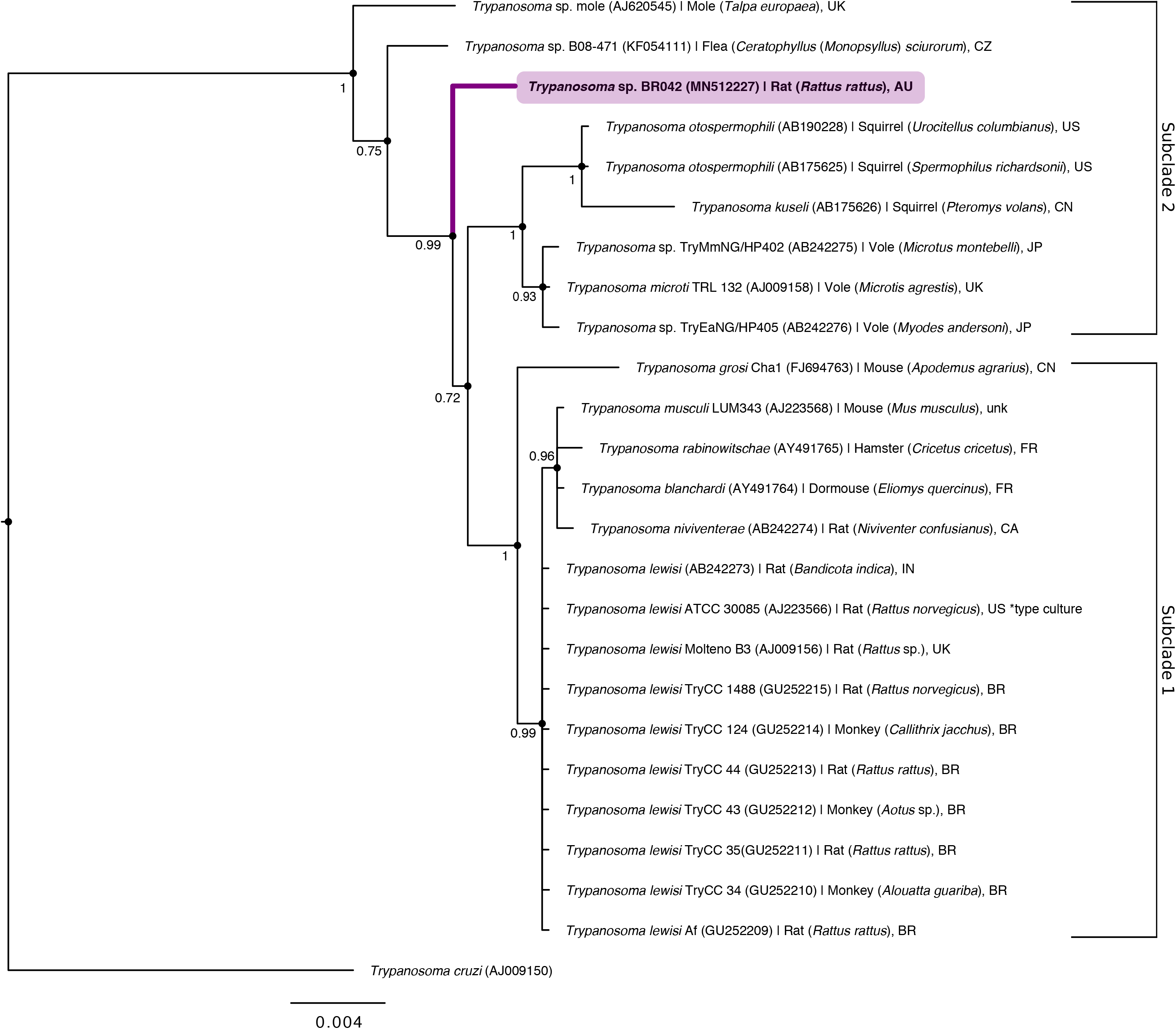
Bayesian phylogenetic reconstruction of *Trypanosoma lewisi* clade using 1,627 bp fragment of 18S rRNA locus based on HKY + G substitution model. Node labels represent posterior probabilities, number of substitutions per nucleotide position are represented by the scale bar. *Trypanosoma cruzi* (AJ009150) was used as group out. New sequence from the present study is designated in bold. Isolate hosts: European mole (*Talpa europaea*), squirrel flea (*Ceratophyllus* (*Monopsyllus*) *sciurorum*), black rat (*Rattus rattus)*, Columbian ground squirrel (*Urocitellus columbianus*), Japanese grass vole (*Microtus montebelli*), Richardson’s ground squirrel (*Spermophilus richardsonii*), Siberian flying squirrel (*Pteromys volans*) field vole (*Microtus agrestis*), Japanese/Anderson’s red-backed vole (*Myodes andersoni),* striped field mouse (*Apodemus agrarius*), house mouse (*Mus musculus*), European hamster (*Cricetus cricetus*), garden dormouse (*Eliomys quercinus*), Chinese white-bellied rat (*Niviventer confusianus*), greater bandicoot rat (*Bandicota indica*), brown rat (*Rattus norvegicus*), brown howler monkey (*Alouatta guariba*), common marmoset (*Callithrix jacchus*), night monkey (*Aotus* sp.). Country abbreviations; Australia (AU), Brazil (BR), Canada (CA), China (CN), Czech Republic (CZ), France (FR), Indonesia (ID), Japan (JP), United Kingdom (UK), United States of America (US), unknown (unk).

Morphological identification of rodent trypanosomes in Australia, attributed to *T. lewisi*, was first made by T. L. Bancroft in 1888 from black rats in Brisbane (Mackerras 1959), with subsequent records by various scientists who confirmed the presence of this parasite in; Brisbane by Pound (1905), in Perth by Cleland (1906, 1908), and in Sydney by Johnston (1909) (cited by Mackerras 1959). *Trypanosoma lewisi* was first identified in native Australian fauna by Mackerras (1958). Morphological detection of the parasite has been made from the bush rat (*Rattus fuscipes*; Queensland) and the water rat (*Hydromys chrysogaster*; Queensland) (Mackerras 1958, 1959). More recently, molecular reports of *Trypanosoma* species from the *T. lewisi* clade have been made from native wildlife in Western Australia, including two bush rats (*Rattus fuscipes*), a dibbler (*Parantechinus apicalis*) and an ash-grey mouse (*Pseudomys albocinereus*) (Averis et al. 2009). Interestingly, despite sampling from 371 native mammals, 19 different species and 14 sites, detection of *T. lewisi*-like species was confined only to mammals from Fitzgerald River in the south-west of Australia. The identification of *T. lewisi*-like spp. by Averis et al. (2009) was limited by the short size of the 18S rRNA locus analysed (444 bp). As demonstrated in the present study, across a short region of the 18S rRNA locus, trypanosomes within the *T. lewisi* clade can share a high sequence similarity (Fig 1b), however upon more robust analysis of a longer fragment it is evident that sequences within the *T. lewisi* clade form distinct groups. Additional genetic information (e.g. glycosomal glyceraldehyde-3-phosphate dehydrogenase (*gGAPDH*)) also assists in determining the phylogenetic relationships of these closely related species. The rabbit trypanosome (*Trypanosoma nabiasi*), which also falls within the *T. lewisi*-clade, has been identified from Australian rabbits and their associated fleas (*Spilopsyllus cuniculi*) in New South Wales and Victoria (Hamilton et al. 2005).

Christmas Island is an external Australian Territory located in the Indian Ocean, south of Indonesia and was once home to endemic populations of *Rattus macleari* and *Rattus nativitatis*. The introduction of black rats and their associated trypanosomes to regions previously free of these species has long been considered responsible for the extinction of two native rat species, a hypothesis that dates back to the time of the extinction events in the early 1900’s by parasitologist H.E. Durham (Durham 1908). Recent research has confirmed Durham’s initial reports and concluded that the rapid decline and extinction of the two endemic rat species was correctly attributed to infections with *T. lewisi* (Wyatt et al. 2008). A review of historical records demonstrated a rapid extinction event following the arrival of black rats on the island in September 1900 and an absence of native rat sightings by October 1904 (Green 2014). While there is strong support for the placement of the trypanosome species responsible within the *T. lewisi* clade, the nature of the ancient DNA study by Wyatt et al. (2008) using museum specimens meant that only a short fragment of the 18S rRNA gene was amplified. As such, differentiation within the *T. lewisi* clade is difficult in this case. A recent study by Dybing et al. (2016) investigated the presence of *Trypanosoma* and *Leishmania* spp. from feral cats (*Felis catus*) and black rats (*R. rattus*) on Christmas Island. Through molecular analysis of spleen samples, the study did not detect any *Trypanosoma* or *Leishmania* species. In addition, the same study reported an absence of these parasites from feral cat samples from Dirk Hartog Island and sites from south-west Western Australia.

North Head is situated on the northern side of Sydney harbour and is dominated by Eastern Suburbs banksia scrub, a declared endangered ecological community (Environment Protection and Biodiversity Conservation Act 1999). In addition to being home to endangered populations of long-nosed bandicoots (*Perameles nasuta*) and little penguins (*Eudyptula minor*), reintroductions of native fauna species, such as bush rats (*Rattus fuscipes*), eastern pygmy possums (*Cercartetus nanus*) and brown antechinus (*Antechinus stuartii*), have also been carried out at North Head by the Australian Wildlife Conservancy. While there is no evidence of spill-over of trypanosomes within the *T. lewisi* clade to native species to-date, ongoing monitoring of populations is encouraged given the historical significance of this parasite with respect to native animal declines (Wyatt et al. 2008; Green 2014).

In addition to trypanosomes, black rats may act as reservoirs for many other sources of infectious agents (Banks and Hughes 2012). Additional information regarding the presence, distribution and diversity of pathogens harboured by black rats in Australia is critical to understanding pathogen spill-over dynamics (Becker et al. 2019). Future research encompassing both morphological and molecular techniques is ongoing by the authors. Collection of ectoparasites, blood, and tissue samples from both native and introduced wildlife will likely continue to shed light on the diversity and distribution of vector-borne microorganisms impacting wildlife, domestic animals and humans.

## Acknowledgements

This study was part-funded by the Australian Research Council (LP160100200), Bayer HealthCare (Germany) and Bayer Australia. S.L.E. is supported by an Australian Government Research Training Program (RTP) Scholarship, C.L.T is supported by a scholarship from the Northern Beaches Council. This project was also part supported by The Holsworth Wildlife Research Endowment & The Ecological Society of Australia (awarded to S.L.E) and the Paddy Pallin Science Grant from The Royal Zoological Society (awarded to C.L.T). We thank Jenna Bytheway, Dr Henry Lydecker and the Australian Wildlife Conservancy ecologists Dr. Viyanna Leo and Mareshell Wauchope for their invaluable assistance in the field.

## Ethical statement

The sampling was conducted under Murdoch University Animal Ethics Committee permit number R3026/18 and University of Sydney Animal Ethics Committee Permit number 2018/1429.

